# NDE1 Localizes to the Subdistal Appendages to Maintain Centrosome Integrity and Microtubule Organization

**DOI:** 10.64898/2026.07.01.735914

**Authors:** Vickie Yang, Paula A. Coelho, Sophia M. Slora, David M. Glover

## Abstract

Centrosomes organize microtubules, coordinate ciliogenesis, and support accurate cell division. At the mature mother centriole, distal and subdistal appendages confer specialized functions in ciliary docking, microtubule organization, and intracellular trafficking. Nuclear distribution element 1 (NDE1) is a centrosome-associated regulator of dynein-dependent processes and ciliogenesis, but its nanoscale organization and contribution to centrosome architecture are poorly understood. Here, using expansion microscopy and STED super-resolution imaging, we show that endogenous NDE1 forms a ring-like structure at the subdistal appendages in mouse embryonic fibroblasts and human RPE-1 cells. NDE1 occupies an intermediate radial position between the more centriole-proximal CEP128 layer and the more peripheral NIN (ninein) layer. Depletion of ODF2 or CEP128 reduces centrosomal NDE1, whereas CEP170 depletion has little effect. Conversely, NDE1 depletion impairs NIN centrosomal enrichment, positioning NDE1 downstream of ODF2 and CEP128 and upstream of NIN within the subdistal appendage assembly hierarchy. NDE1 depletion compromises centrosome integrity, reduces the centrosomal enrichment of core centriolar proteins, increases the separation between paired centrioles, and generates ectopic foci containing multiple centriolar markers. Loss of NDE1 also disrupts pericentriolar material organization and impairs the establishment of focused, centrosome-associated microtubule arrays. Together, our findings establish NDE1 as a subdistal appendage-associated factor that supports centrosome architecture and microtubule-organizing activity. More broadly, they support an emerging view of subdistal appendages as a molecularly layered platform in which distinct but cooperating components connect mother centriole maturation to microtubule organization, ciliary regulation, and intracellular trafficking.

## Introduction

Centrosomes are the principal microtubule-organizing centers of most animal cells. Each centrosome typically comprises a pair of centrioles (mother and daughter) surrounded by the microtubule-nucleating pericentriolar material (PCM). Centrosomes organize interphase and mitotic microtubules to ensure cell shape and polarity, mitotic spindle assembly, and accurate chromosome segregation. When a primary cilium is assembled, the mature mother centriole also serves as its basal body. Centrosome architecture, number, and maturation must therefore be coordinated with both cell-cycle progression and ciliogenesis^1^.

The mature mother centrioles of mammalian cells are distinguished by distal and subdistal appendages. Distal appendages mediate ciliary vesicle docking and are required for the initiation of ciliogenesis, whereas subdistal appendages contribute to microtubule organization and have also been implicated in intracellular trafficking and ciliary regulation. Subdistal appendages have a molecularly ordered architecture in which ODF2, CEP128, CNTRL (centriolin), NIN (ninein) and CEP170 occupy distinct radial and axial positions. Their assembly is interdependent: ODF2 and CEP128 function relatively early and support the recruitment or organization of more peripheral appendage-associated factors^2–6^. The acquisition of these appendages is a hallmark of centriole maturation, through which a newly formed daughter centriole progressively acquires mother-specific structures and full microtubule-organizing and basal-body competence^7^.

Nuclear distribution element 1 (NDE1) is a multifunctional centrosome-associated protein^8,9^. NDE1 interacts with dynein regulators, including LIS1, and contributes to mitotic progression, spindle organization, nuclear positioning and migration, and neurodevelopment^10–14^. NDE1 also negatively regulates ciliary length: its depletion promotes ciliary elongation and can delay cell-cycle re-entry^15^, whereas cell-cycle-dependent degradation of NDE1 through the CDK5-FBW7 pathway facilitates ciliogenesis^16^. Consistent with essential functions in the proliferation and migration of neural progenitor cells, pathogenic loss-of-function variants in NDE1 cause severe neurodevelopmental disorders characterized by profound microcephaly and cortical malformations^17^. Despite the established centrosomal functions, the nanoscale organization of NDE1 at the centrosome and its contribution to centriole architecture have remained unclear.

Here, using expansion microscopy and STED super-resolution imaging (ExSTED)^18^, we define the nanoscale organization of NDE1 at the subdistal appendage region. We show that NDE1 forms a ring-like structure at the mother centriole and occupies an intermediate radial position within the ordered subdistal appendage architecture. Centrosomal NDE1 is reduced following depletion of ODF2 or CEP128 but is largely unaffected by CEP170 depletion, whereas NDE1 depletion impairs NIN centrosomal localization, positioning NDE1 within the subdistal appendage recruitment network. NDE1 depletion broadly alters centriolar protein enrichment, increases the separation between paired centrioles, and generates ectopic centriolar-marker-positive foci. Loss of NDE1 also disrupts pericentriolar material organization and compromises the establishment of focused centrosome-associated microtubule arrays. Together, our findings identify NDE1 as a subdistal appendage-associated factor that supports centrosome architecture and microtubule-organizing activity and expands the functional repertoire of the mother centriole subdistal appendage region.

## Results

### NDE1 localizes to the mother centriole subdistal appendage region

NDE1 has previously been detected at centrosomes, but its precise spatial organization within the centrosome has not been established. To define its nanoscale distribution, we examined the localization of endogenous NDE1 using expansion microscopy and STED super-resolution imaging (ExSTED) in mouse embryonic fibroblasts (MEFs; Fig. 1 A, S1 A). In top-down views of interphase mother centrioles, NDE1 formed a ring surrounding the acetylated-tubulin-positive centriole cylinder. In side views, NDE1 localized laterally to the distal, appendage-bearing region of the mother centriole. Additional NDE1 signal was detected throughout the cytoplasm. Although the identity of this cytoplasmic pool was not investigated, we hypothesize it may represent the NDE1 population binding cytoplasmic dynein. A comparable centrosomal distribution was observed in human RPE-1 cells (Fig. S1 B), indicating that this localization is shared between the two mammalian cell types examined.

**Figure 1.**
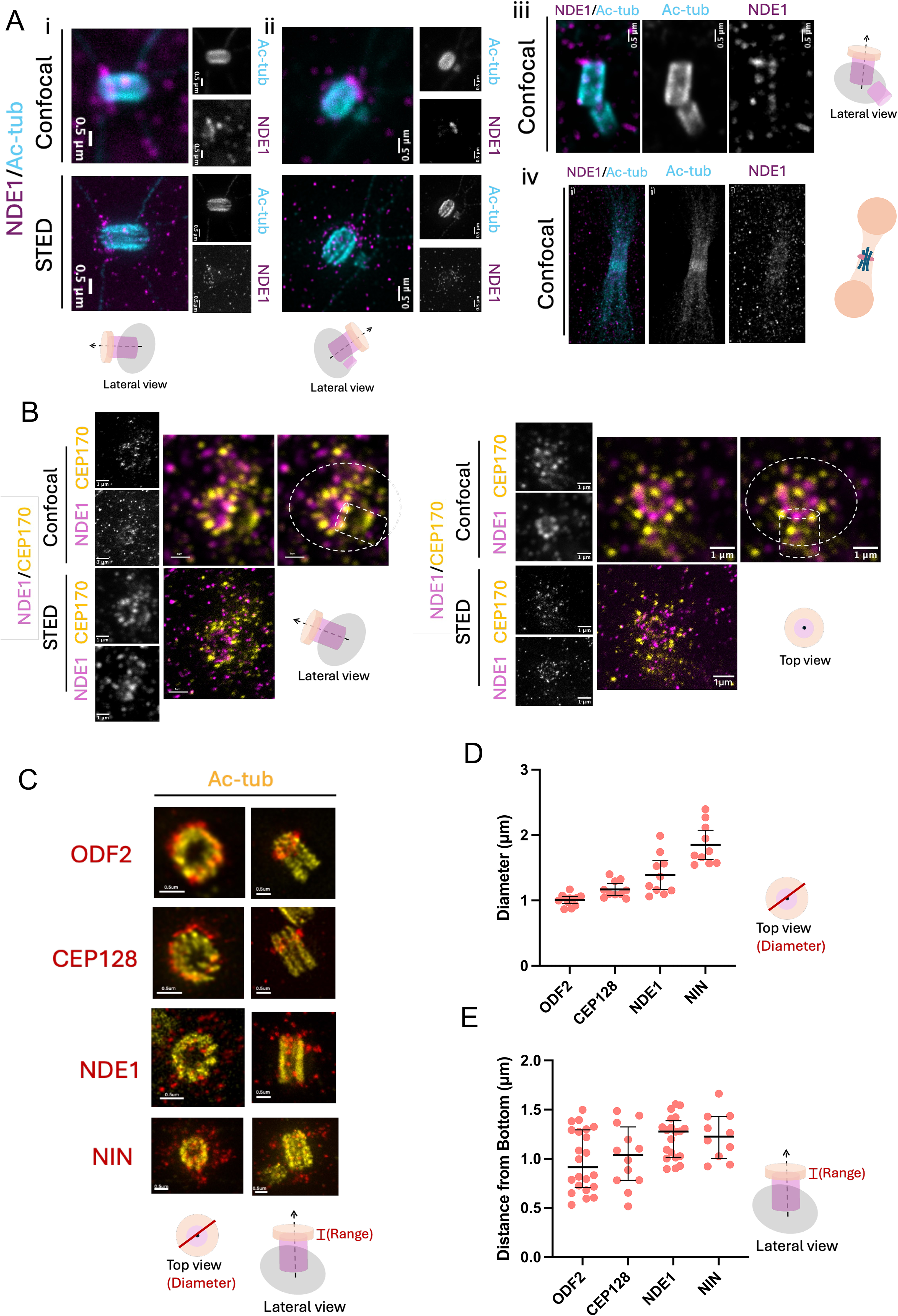
NDE1 localizes to subdistal appendage region of the mother centriole. **(A)** Deconvolved ExM-confocal and ExSTED images showing NDE1 localization at the appendage-bearing region of the mother centriole during interphase in MEFs. Representative G1 and S/G2 centrosomes are shown in panels i and ii, respectively. The centriole cylinder is marked by acetylated tubulin (Ac-tub, cyan), and NDE1 is shown in magenta. During mitosis, NDE1 staining at the appendage region is reduced (iii), and NDE1 is also detected at the contractile ring (iv). Schematics below the images indicate the corresponding centriole orientation. Scale bars, 0.5 µm in i-iii and 1 µm in iv. **(B)** ExM-confocal and ExSTED images showing the spatial relationship between NDE1 and the subdistal appendage protein CEP170 at the mother centriole in RPE-1 cells. NDE1 is shown in magenta, and CEP170 is shown in yellow. Lateral and top-down views are shown in the left and right panels, respectively. Dashed circles outline the appendage-associated domain, and dashed rectangles indicate the approximate position and orientation of the centriole cylinder. Schematics indicate the centrosome orientations represented in the corresponding images. Scale bars, 1 µm. **(C)** Representative ExSTED images comparing the radial and axial localization of ODF2, CEP128, NDE1, and ninein (NIN) at the mother centriole in MEFs. Top-down and lateral views are shown on the left and right, respectively. The indicated subdistal appendage protein is shown in red, and Ac-tub is shown in yellow. Schematics indicate the centriole orientations represented in the corresponding images. Scale bars, 0.5 µm. **(D)** Quantification of ODF2, CEP128, NDE1, and NIN ring diameters from top-down ExSTED images. Each point represents one mother centriole. Black bars indicate the mean and 95% confidence interval. Exact sample sizes: ODF2, n = 12; CEP128, n = 10; NDE1, n = 10; and NIN, n = 10 from 3 independent experiments. **(E)** Quantification of the axial positions of ODF2, CEP128, NDE1, and NIN relative to the Ac-tub-marked centriole base in lateral ExSTED images. Each point represents one mother centriole measurement. Black bars indicate the median and interquartile range. Exact sample sizes: ODF2, n = 22; CEP128, n = 12; NDE1, n = 20; and NIN, n = 10 from 3 independent experiments.

Across the interphase stages examined, NDE1 remained enriched at the mother centriole and displayed an appendage-associated distribution. Weaker NDE1 signal was also occasionally detected near the proximal region of the centriole. During cytokinesis, a fraction of NDE1 redistributed to the cytokinetic apparatus, whereas centrosomal signal appeared reduced (Fig. 1 A, S1 A). This cell-cycle-dependent pattern resembles the mitotic remodeling reported for peripheral subdistal appendage proteins, including centriolin, NIN and CEP170^3,6^, and raises the possibility that the NDE1-associated subdistal appendage domain is reorganized during mitosis.

To determine whether NDE1 occupies the subdistal appendage region, we co-stained NDE1 with the established subdistal appendage marker CEP170. In both top-down and side views, NDE1 and CEP170 occupied overlapping spatial domains around the mother centriole (Fig. 1 B, S1 C). Although spatial overlap at the resolution achieved here does not demonstrate a direct molecular interaction, these observations strongly suggest localization of NDE1 to the subdistal appendage region.

We next compared the radial organization of NDE1 with that of other established subdistal appendage components. STED imaging resolved ring-like distributions of ODF2, CEP128, NDE1, and NIN around the mother centriole. Quantification of ring diameter revealed a radially ordered arrangement, with the ODF2 ring having the smallest diameter, followed by CEP128, NDE1, and NIN (Fig. 1, C and D, S1 D). NDE1 therefore occupies an intermediate radial position within the subdistal appendage architecture, with an average ring diameter of 1.388 ± 0.310 μm (mean ± SD, n = 10) in the 4x expanded sample, positioned between the more centriole-proximal CEP128 layer and the more peripheral NIN layer.

We further mapped the axial distributions of these proteins along the proximal-distal centriole axis. Longitudinally oriented mother centrioles were aligned along this axis, and the acetylated-tubulin-positive centriole cylinder was used as a common spatial reference. ODF2 and CEP128 occupied comparatively broad axial domains centered toward the proximal portion of the appendage-bearing region. By contrast, NDE1 and NIN displayed narrower distributions centered at more distal axial positions, with NDE1 located at an average distance of 1.218 ± 0.215 μm (mean ± SD, n = 20) from the proximal end of the centriole in the expanded sample (Fig. 1, C and E, S1 E). Together, the radial and axial measurements position NDE1 within a distinct nanoscale domain of the mother-centriole subdistal appendage region.

### ODF2 and CEP128 support centrosomal accumulation of NDE1

Subdistal appendage proteins are recruited through a hierarchical pathway. We therefore asked whether established appendage components were required for centrosomal localization of NDE1. Depletion of ODF2 reduced NDE1 centrosomal localization, consistent with ODF2 acting as a core upstream scaffold for appendage assembly. Similarly, CEP128 depletion markedly impaired NDE1’s enrichment to the subdistal appendage region. In contrast, depletion of CEP170 did not substantially reduce centrosomal NDE1 localization (Fig. 2 A and B). These results suggest that accumulation of NDE1 at the mother centriole requires ODF2 and CEP128 but is largely independent of CEP170. Conversely, ExSTED indicated that NDE1 depletion reduced centrosomal NIN signal, in accord with NDE1 contributing to the assembly or maintenance of more peripheral subdistal appendage components.

**Figure 2.**
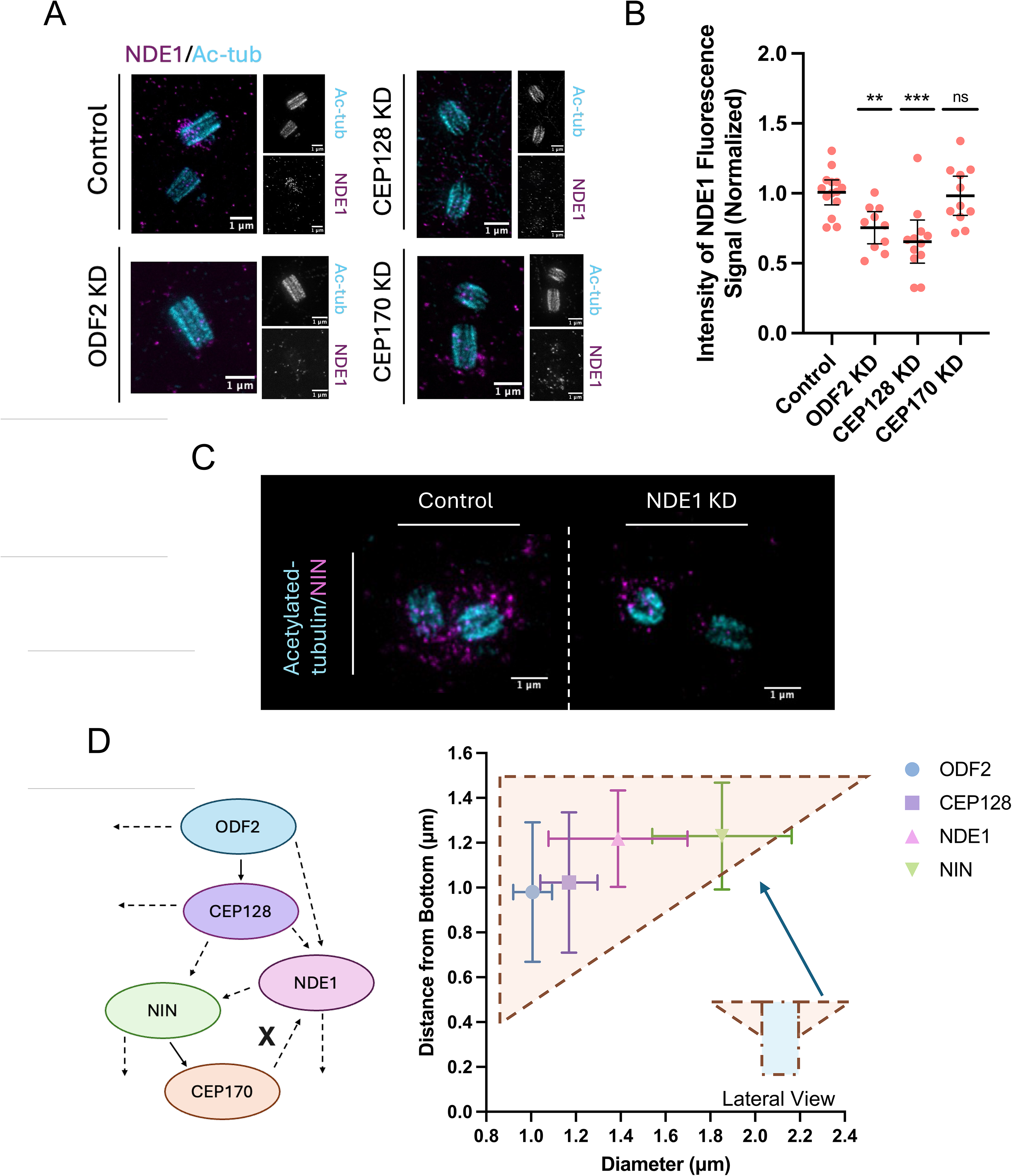
ODF2 and CEP128 support centrosomal accumulation of NDE1 in MEFs. **(A)** Representative ExSTED images of NDE1 staining in control, ODF2-depleted, CEP128-depleted, and CEP170-depleted cells (MEFs). The centriole cylinder is marked by acetylated tubulin (Ac-tub, cyan), and NDE1 is shown in magenta. Scale bar, 1 µm. **(B)** Quantification of centrosomal NDE1 fluorescence intensity in the indicated conditions from ExSTED images. Each point represents one mother centriole. Black bars indicate mean with 95% confidence interval. Statistical significance was determined by one-way ANOVA followed by Dunnett’s multiple-comparisons test comparing each depletion condition with the control; ***P < 0.001; **P < 0.01; ns, not significant. Exact sample sizes: Control, n = 14; ODF2 KD, n = 10; CEP128 KD, n = 12; and CEP170 KD, n = 11 from 3 independent experiments. **(C)** Representative ExSTED images showing NIN centrosomal enrichment in control and NDE1-depleted cells. The centriole cylinder is marked by acetylated tubulin (Ac-tub, cyan), and NIN is shown in magenta. Scale bar, 1 µm. **(D)** Schematic model of the subdistal appendage assembly (left). Solid lines indicate previously reported biochemical interactions, whereas dashed lines indicate epistasis or recruitment relationships inferred from depletion experiments. The cross indicates that centrosomal recruitment of NDE1 is largely independent of CEP170. Two-dimensional spatial mapping of ODF2, CEP128, NDE1, and NIN within the subdistal appendage in MEFs (right). Mean protein positions are plotted according to the measured diameter (x axis) and distance from the bottom of the mother centriole (y axis). Horizontal and vertical error bars indicate SD. The overlaid schematic depicts the approximate lateral geometry of the subdistal appendage and its relationship to the centriole.

Together with the radial mapping, this depletion analysis positions NDE1 downstream of ODF2 and CEP128 in subdistal appendage assembly, but upstream of NIN, while remaining largely independent of CEP170 (Fig. 2 C). We observed a comparable outcome in human RPE-1 cells (Fig. S2 A-C).

### NDE1 depletion compromises centriole architecture and generates ectopic centriolar-marker-positive foci

Having established that NDE1 localizes to the subdistal appendage region, we next asked whether NDE1 contributes to centriole organization. Following NDE1 depletion, both MEFs and RPE-1 cells displayed abnormal centriole structures. ExSTED revealed reduced centriolar fluorescence intensities of CEP135 and acetylated tubulin, accompanied by distorted centriole morphology (Fig. 3 A). Similarly, the centriolar accumulation of CPAP and CEP152 was also reduced, together with alterations in the organization of their associated centriole structures (Fig. 3 C-F, S3 A-D). Because these proteins occupy distinct structural and regulatory domains of the centriole^19,20^, their coordinated reduction indicates that NDE1 depletion broadly disrupts centriole organization. These changes may reflect altered centriolar protein recruitment or retention.

**Figure 3.**
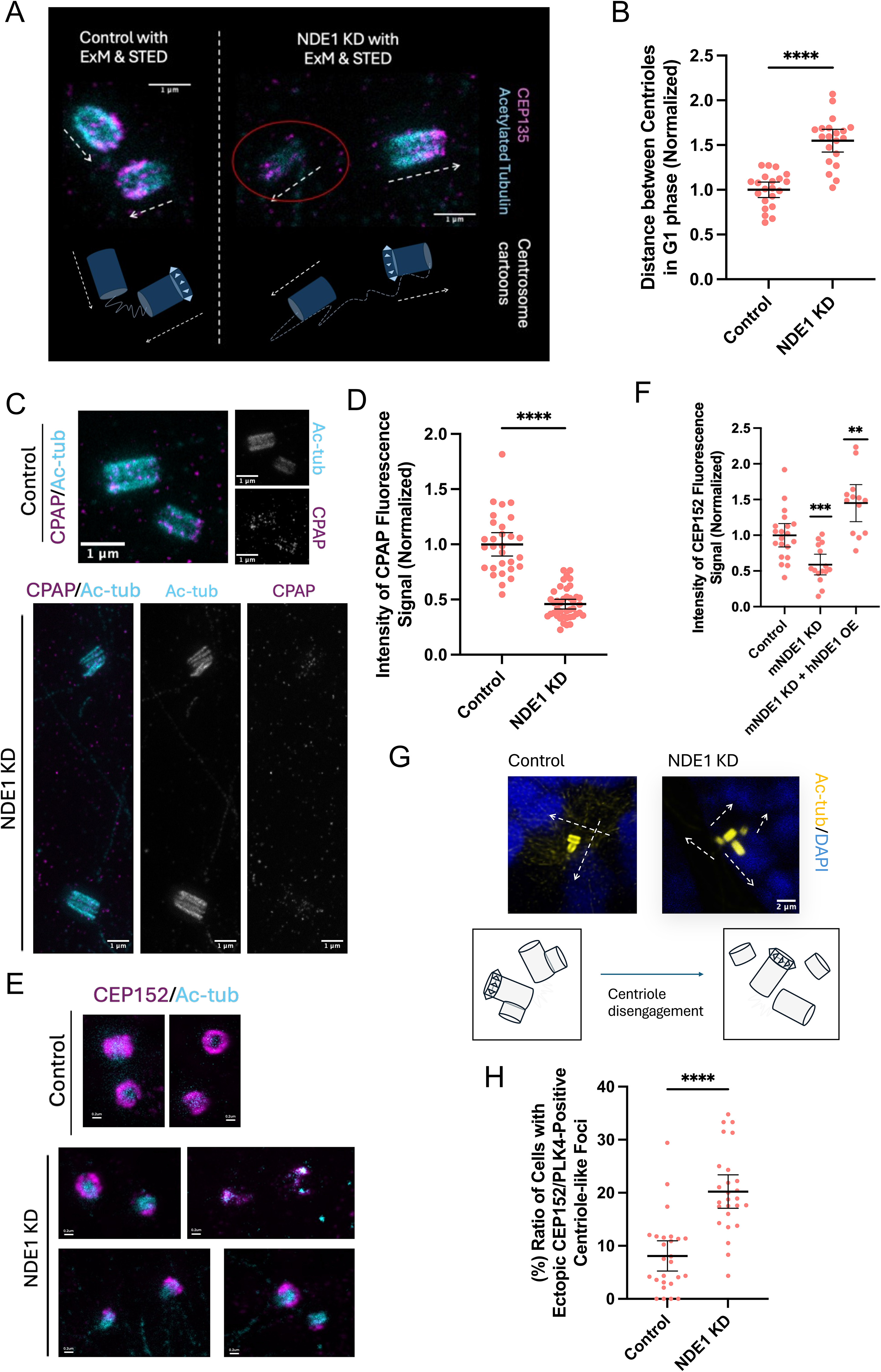
NDE1 depletion compromises centriole architecture and generates ectopic centriolar-marker-positive foci. **(A)** Representative ExSTED images showing reduced CEP135 staining in NDE1-depleted MEFs. NDE1 depletion is associated with increased distance between paired centrioles and abnormal centriole morphology. The centriole cylinder is marked by acetylated tubulin (Ac-tub, cyan), and CEP135 is shown in magenta. White arrows and schematics below the images indicate centriole orientation. Red circles highlight abnormal centriole structures. Scale bars, 1 µm. **(B)** Quantification of the distance between paired centrioles in G1-phase MEFs under control and NDE1-depleted conditions from confocal images. Each point represents one individual cell. Black bars indicate the mean with 95% confidence interval. Statistical significance was determined by two-tailed unpaired t test; ****P < 0.0001. Exact sample sizes: Control, n = 21; and NDE1 KD, n = 20 from 3 independent experiments. **(C)** Representative ExSTED images showing reduced CPAP staining in NDE1-depleted MEFs. NDE1 depletion is associated with increased distance between paired centrioles and abnormal centriole morphology. The centriole cylinder is marked by Ac-tub, and CPAP is shown in magenta. Scale bars, 1 µm. **(D)** Quantification of centriolar CPAP fluorescence intensity in control and NDE1-depleted MEFs. Each point represents one individual cell. Black bars indicate the mean with 95% confidence interval. Statistical significance was determined by two-tailed unpaired t test; ****P < 0.0001. Exact sample sizes: Control, n = 29; and NDE1 KD, n = 41 from 3 independent experiments. **(E)** Representative STED images showing reduced CEP152 staining in NDE1-depleted MEFs. NDE1 depletion is associated with increased distance between paired centrioles and abnormal centriole morphology. The centriole cylinder is marked by Ac-tub, and CEP152 is shown in magenta. Scale bars, 0.2 µm. **(F)** Quantification of centriolar CEP152 fluorescence intensity in control and NDE1-depleted MEFs. Each point represents one individual cell. Black bars indicate the mean with 95% confidence interval. Statistical significance was determined by two-tailed unpaired t test; ***P < 0.001; **P < 0.01. Exact sample sizes: Control, n = 20; mNDE1 KD, n = 15; and mNDE1 KD+hNDE1 OE, n = 13 from 3 independent experiments. **(G)** Representative ExM images showing altered centriole-pair organization in NDE1-depleted MEFs. Loss of orthogonality and increased distance between paired centrioles are observed. The centriole cylinder is marked by Ac-tub in yellow, and nuclei are stained with DAPI in blue. Schematics below the images indicate centriole orientation. Scale bars, 2 µm. **(H)** Quantification of the percentage of cells with ectopic CEP152/PLK4-positive centriole-like foci in control and NDE1-depleted MEFs. Each point represents one individual field. Black bars indicate the mean with 95% confidence interval. Statistical significance was determined by two-tailed unpaired t test; ****P < 0.0001. Exact sample sizes: Control, n = 26; and NDE1 KD, n = 26 from 3 independent experiments.

NDE1-depleted cells also displayed increased separation between paired centrioles and, in some cases, premature disintegration of procentrioles (Fig. 3 B, G, S3 A-D), indicating that NDE1 contributes to the normal structural and spatial organization of centriole pairs. Although altered centriole spacing has also been reported following disruption of some other subdistal appendage proteins^6^, its underlying mechanism in NDE1-depleted cells remains to be determined.

Interestingly, NDE1-depleted cells contained ectopic puncta positive for multiple centriolar markers (Fig. 3 H). A subset of these foci was associated with local microtubule asters, suggesting that some retained microtubule-organizing capacity. Collectively, these foci likely represent a heterogeneous population of centriole-related structures generated upon NDE1 depletion, including structurally compromised centrioles, prematurely separated centrioles, incompletely assembled centriole-related structures, or centriole-derived fragments that retain centrosomal components.

Together, these findings indicate that NDE1 is required to generate or maintain normal centriole protein organization and centrosome architecture in both MEFs and RPE-1 cells. The accumulation of ectopic centriolar-marker-positive foci, some of which retain apparent microtubule-organizing capacity, further suggests that NDE1 depletion alters the relationship between centriole structure, centrosome organization, and the cellular recognition of centrosome number.

### NDE1 supports PCM organization and centrosome-focused microtubule growth

Because subdistal appendages contribute to the anchoring of interphase microtubules at the centrosome^3,4^, we asked whether NDE1 depletion altered the centrosomal microtubule-organizing machinery. Our results showed that centrosomal γ-tubulin signal was relatively unchanged, whereas pericentrin was more strongly diminished (Fig. 4 A)^21^. These changes indicate that NDE1 depletion perturbs PCM organization and may thereby compromise centrosomal microtubule organization. We next examined microtubule organization in interphase and mitotic cells. In interphase, NDE1-depleted cells displayed less focused centrosome-centered microtubule arrays. In mitotic cells, the centrosome-proximal EB1 signal was also reduced (Fig. 4 B-D)^22^. Comparable findings were made in human RPE-1 cells depleted of NDE1 (Fig. S4 A-F).

**Figure 4.**
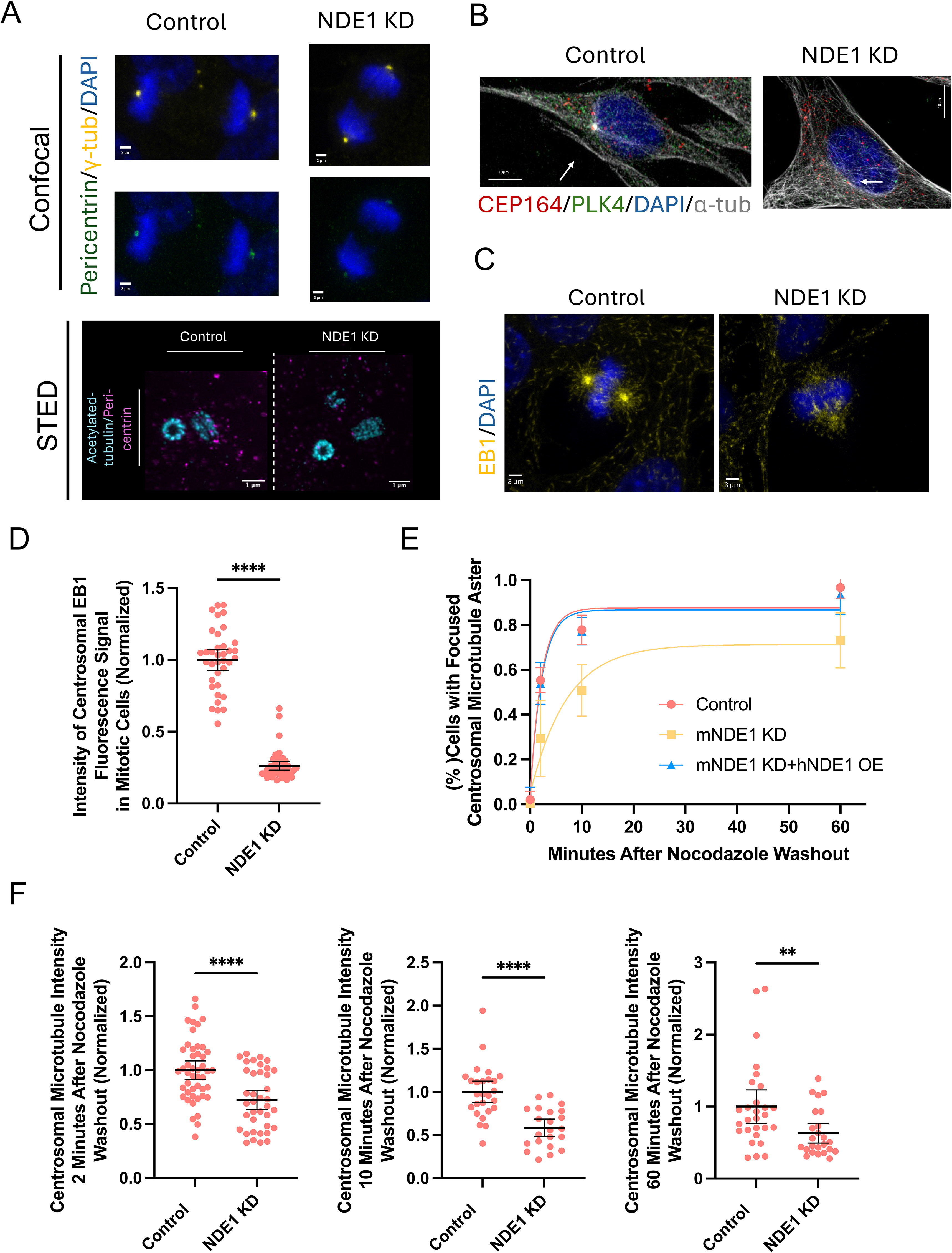
NDE1 supports PCM organization and centrosome-focused microtubule growth in MEFs. **(A)** Representative confocal images showing reduced centrosomal pericentrin staining in NDE1-depleted MEFs (Upper). γ-tubulin, which marks the centrosome and is comparatively less affected by NDE1 depletion, is shown in yellow; pericentrin is shown in green; and DNA is stained with DAPI in blue. Scale bars, 3 µm. Representative ExSTED images showing reduced centrosomal pericentrin staining and an ectopic centriole-like fragment in NDE1-depleted MEFs (Lower). The centriole cylinder is marked by acetylated tubulin (Ac-tub, cyan), and pericentrin is shown in magenta. Scale bars, 1 µm. **(B)** Representative confocal images showing reduced centrosome-focused microtubule organization in interphase NDE1-depleted MEFs. Microtubules are shown in gray, CEP164 in red, PLK4 in green, and nuclei are stained with DAPI in blue. Centrioles are defined by CEP164/PLK4-positive foci. Scale bars, 10 µm. **(C)** Representative confocal images showing reduced centrosome-focused microtubule asters in mitotic NDE1-depleted MEFs. EB1-positive microtubule plus ends are shown in yellow, and DNA is stained with DAPI in blue. Scale bars, 3 µm. **(D)** Quantification of centrosome-associated EB1 fluorescence intensity during mitosis in control and NDE1-depleted MEFs. Each dot represents the EB1 fluorescence intensity associated with an individual centrosome. Black bars indicate the mean with 95% confidence interval. Statistical significance was determined by two-tailed unpaired t test for comparison with the control condition; ****P < 0.0001. Exact sample sizes: Control, n = 36; and NDE1 KD, n = 44 from 3 independent experiments. **(E)** Time course of centrosomal microtubule aster recovery following nocodazole washout, quantified as the fraction of cells displaying a focused centrosomal microtubule aster in control (red), NDE1-depleted (yellow), and NDE1-depleted MEFs expressing human NDE1 (hNDE1; blue) following nocodazole washout. Curves were fitted using nonlinear regression with a one-phase association model. Data are shown as mean ± SD. At 0, 2, 10, and 60 min after nocodazole washout, exact sample sizes were: Control, n = 7, 6, 9, and 8; mNDE1 KD, n = 7, 6, 7, and 6; and NDE1 KD + hNDE1 OE, n = 7, 6, 7, and 8, respectively. **(F)** Quantification of centrosomal α-tubulin intensity at 2, 10, and 60 minutes after nocodazole washout in control and NDE1-depleted MEFs. Each dot represents the centrosome-associated α-tubulin fluorescence intensity of an individual centrosome. Black bars indicate the mean with 95% confidence interval. Statistical significance was determined by two-tailed unpaired t test for comparison to the corresponding control condition; ****P < 0.0001, **P < 0.01. At 2 minutes after nocodazole washout, exact sample sizes: Control, n = 46; and NDE1 KD, n = 38 from 3 independent experiments. At 10 minutes after nocodazole washout, exact sample sizes: Control, n = 26; and NDE1 KD, n = 22 from 3 independent experiments. At 60 minutes after nocodazole washout, exact sample sizes: Control, n = 28; and NDE1 KD, n = 24 from 3 independent experiments.

We further assessed centrosomal microtubule organization using a nocodazole washout assay. NDE1 depletion reduced the fraction of cells displaying a focused centrosomal microtubule aster as early as 2 minutes after washout, when microtubules begin to regrow from the centrosomal region, and this defect persisted at later time points (10–60 minutes), indicating impaired re-establishment of centrosome-focused microtubule organization (Fig. 4 E; Fig. S4 G). Consistently, centrosome-associated microtubule intensity remained lower in NDE1-depleted cells throughout the regrowth time course (Fig. 4 F). Together, these findings support a role for NDE1 in PCM organization and the efficient establishment of centrosome-centered microtubule arrays.

We next asked whether the disruption of centrosome-focused microtubule organization following NDE1 depletion was accompanied by changes in intracellular membrane compartments. We focused on the autophagic pathway, whose compartments undergo microtubule-dependent transport and maturation^23–25^. NDE1 depletion, however, revealed no significant effect on bulk autophagic flux in either MEFs or RPE-1 cells (Fig. S5 A-E; Fig. S6 A-G). Consistently, SQSTM1/p62 levels were only modestly increased and continued to accumulate following BafA1 treatment^26^. By contrast, immunofluorescence revealed a prominent increase in puncta-associated LC3B signal following NDE1 depletion (Fig. S5 F), which was restored toward control levels by expression of human NDE1 (Fig. S5 G). Similarly, NDE1 depletion significantly increased autophagosome number without significantly altering autolysosome number (Fig. S5 H and I). LC3B-positive structures also remained abundant in NDE1-depleted cells following prolonged BafA1 treatment (Fig. S6 H), and mCherry-LAMP1-positive compartments were not significantly increased (Fig. S6 I). Thus, the increase in autophagosome number was not accompanied by a detectable change in bulk autophagic flux or an increase in LAMP1-positive compartments (Fig. S7).

## Discussion

Although NDE1 has been previously implicated in centrosome function, dynein-dependent processes, and ciliary regulation, its nanoscale organization at the centrosome has remained unclear. Here, we show that endogenous NDE1 forms an annular region around the centriole in the position of the subdistal appendages and occupies an intermediate radial position between the more centriole-proximal CEP128 layer and the more peripheral NIN layer. This spatial organization places NDE1 near the interface between the inner appendage scaffold and peripheral domains associated with microtubules and motor-related machinery, consistent with the established functional relationship between NDE1 and cytoplasmic dynein regulators. Our findings suggest a model whereby NDE1 is recruited to the subdistal appendages downstream of ODF2 and CEP128, where it maintains centrosome structural integrity, supports centrosome-focused microtubule growth, and may affect microtubule-dependent compartment trafficking (Fig. 5).

**Figure 5.**
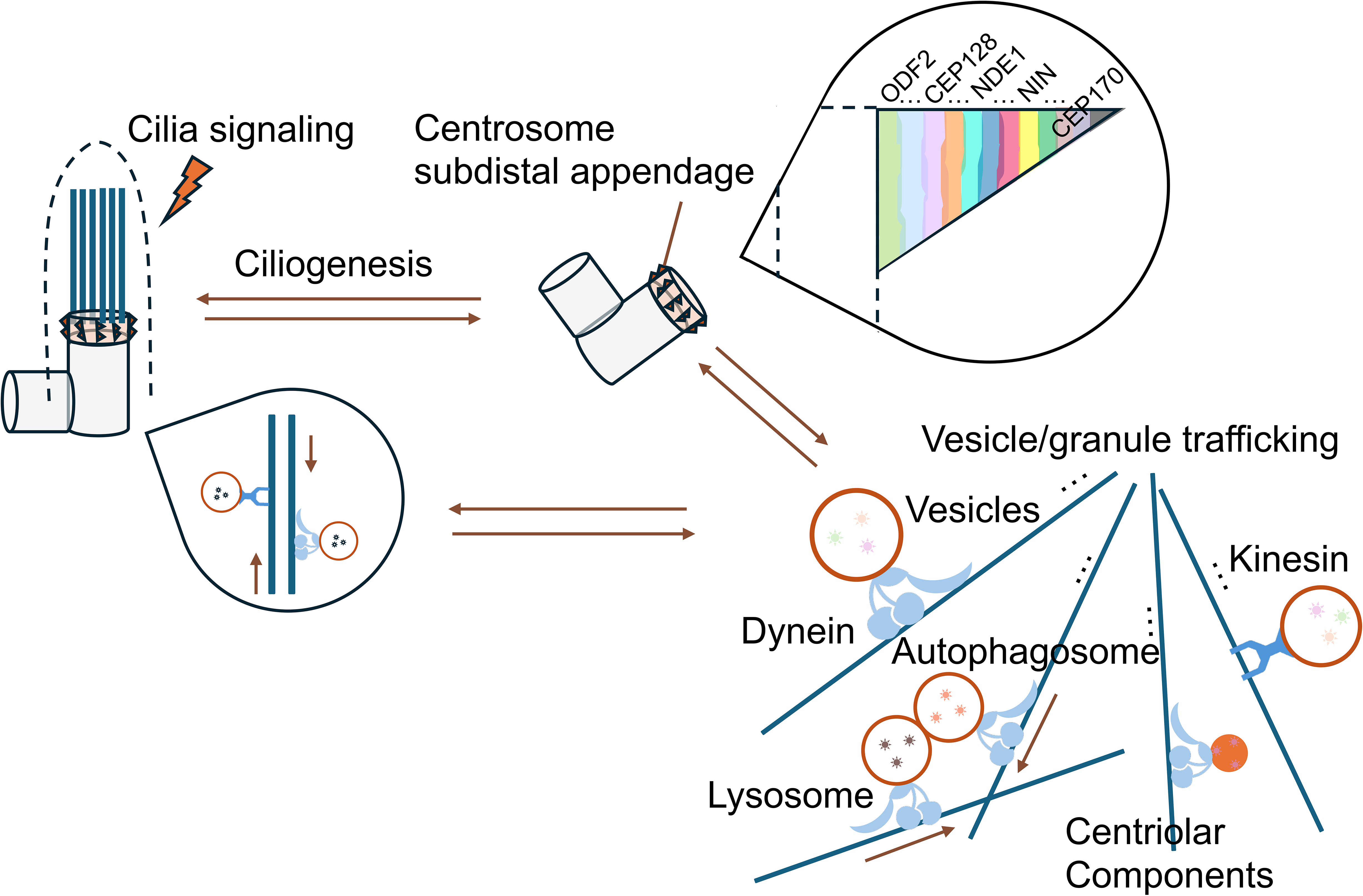
Proposed model for NDE1 function at the subdistal appendage region. NDE1 localizes to the subdistal appendage region of the mother centriole, within an ordered appendage-associated protein network that includes ODF2, CEP128, NIN, and CEP170. Our data support a model in which ODF2 and CEP128 promote centrosomal accumulation of NDE1, and NDE1 in turn contributes to centrosome structural integrity, PCM organization, and centrosome-focused microtubule organization. Through its association with the subdistal appendage region and dynein-related machinery, NDE1 may also help coordinate microtubule-dependent trafficking of vesicles, granules, autophagosomes, lysosomes, and centriolar components near the centrosome. Together with its previously described role as a negative regulator of ciliogenesis, this model places NDE1 at an interface between subdistal appendage organization, microtubule-dependent trafficking, centrosome integrity, and ciliary regulation.

The distinct appearances of the appendage-associated rings in our super-resolution images may reflect differences in the organization of these molecular domains. ODF2 and CEP128 formed comparatively compact and regularly delineated rings, whereas NDE1 displayed a broader and less sharply defined distribution, together with additional signal near the proximal regions. NIN formed a larger and more heterogeneous ring and showed substantial signal near the proximal or cohesion-associated region. One interpretation is that the inner appendage layers form a relatively stable and constrained structural scaffold, whereas proteins positioned further from the centriole wall occupy more spatially heterogeneous domains that engage microtubules, motors, or transported material. NDE1 may reside near the boundary between these structural and dynamic layers. Differences in apparent ring morphology can also result from protein abundance and their relative affinity to the subdistal appendage region.

Centrosomal NDE1 was reduced following depletion of ODF2 or CEP128 but was largely retained after CEP170 depletion. Conversely, NDE1 depletion impaired centrosomal recruitment of NIN, suggesting that NDE1 contributes to the organization or stability of the outer subdistal appendage domain. Together, these findings are consistent with a hierarchical organization in which NDE1 depends on the ODF2-CEP128 scaffold and, in turn, is required for efficient recruitment or maintenance of NIN at subdistal appendages. Nevertheless, the depletion experiments establish architectural dependence rather than direct molecular recruitment. Loss of ODF2 or CEP128 could reduce NDE1 indirectly by disrupting appendage assembly and stability. Direct interaction mapping and localization-specific rescue experiments will therefore be required to determine how NDE1 is incorporated into subdistal appendages and whether more than one recruitment pathway contributes to its centrosomal localization.

NDE1 depletion affected proteins distributed across several centriolar domains: the centriolar microtubule walls marked by acetylated tubulin; the proximal cartwheel-associated architecture associated with CEP135; the centriolar microtubule assembly and elongation machinery marked by CPAP; and the peripheral part of the centriole marked by CEP152, which provides a proximal scaffold for PLK4 recruitment and procentriole initiation. The pleiotropy of cellular phenotypes together with altered centriole morphology suggests that loss of NDE1 produces a broad disturbance of centriole architecture rather than selectively affecting specific subdistal appendage-associated components. That said, NDE1 is required to maintain the normal molecular organization of the centrosome.

Several mechanisms could account for these centriole phenotypes. NDE1 may connect the appendage region to centrosomal microtubules and/or the pericentriolar material to help stabilize centriole organization and maintain the local concentration of centrosomal components. Its link to dynein can expose the centriole microtubules to local dynein-dependent forces. Alternatively, the centriole abnormalities could arise secondarily after prolonged dynein-mediated disruption of microtubule organization, intracellular transport, centrosome positioning, or cell-cycle progression. Knowledge of the timing of these phenotypes relative to the changes in cytoplasmic and centriole microtubules and pericentriolar material will be important for distinguishing primary from secondary effects. We also observed ectopic puncta positive for multiple centriolar markers upon NDE1 knockdown that could represent a heterogeneous population of centriole-related structures. Live tracking of centriole assembly, maturation, and disengagement markers will be required to determine how these structures arise.

The effects of NDE1 depletion on pericentriolar material and microtubule organization further support its role in coordinating the interactions of the mother centriole with the surrounding microtubule-organizing machinery. NDE1-depleted cells displayed a pronounced reduction in centrosomal pericentrin, diminished centrosome-proximal EB1 signal, and less focused interphase microtubule arrays. Consistent with these steady-state defects, microtubule regrowth following nocodazole washout was impaired, indicating a reduced capacity to rapidly re-establish a focused centrosomal microtubule network. Altered pericentriolar material could reduce the generation or stabilization of new microtubules. Moreover, disruption of the subdistal appendage region could impair microtubule anchoring or retention. Defective dynein-associated functions could prevent newly formed microtubules from being efficiently focused toward the centrosome. NDE1 may therefore coordinate several related processes rather than acting exclusively through a single peripheral microtubule-binding mechanism.

Together, these findings support an emerging view of subdistal appendages as multifunctional organizational domains whose activities extend beyond microtubule anchoring to include centrosome positioning, mother-centriole maturation, ciliary regulation, and intracellular trafficking. These processes are closely interconnected: centrosomal microtubules provide tracks for motor-dependent transport, trafficking pathways deliver components to the centrosome and cilium, and ciliary assembly and disassembly must be coordinated with centrosome function and cell-cycle progression. Subdistal appendages are therefore well positioned to couple the maturation and functional state of the mother centriole to the spatial organization of the surrounding cytoplasm. The distinct phenotypes resulting from perturbation of ODF2, CEP128, centriolin, NIN, CEP170, and NDE1 further suggest that these activities are distributed among specialized but cooperating molecular modules. Defining how these modules are assembled and functionally coordinated will be crucial for understanding how the mature mother centriole integrates centrosome architecture, microtubule organization, and cytoplasmic trafficking.

The autophagy-associated phenotype following NDE1 depletion may reflect a broader consequence of disrupted intracellular organization. Although bulk autophagic flux was not detectably reduced, the selective accumulation of autophagosomes suggests an altered balance between autophagic compartments. Given the dependence of autophagosome trafficking and maturation on microtubule-based transport and interactions with lysosomes, disruption of the NDE1-dependent centrosomal microtubule network could contribute to this phenotype. Whether autophagosome accumulation results from altered biogenesis, transport, maturation, or compartment lifetime remains to be determined. In addition, NDE1 depletion was accompanied by a more dispersed cytoplasmic distribution of CEP164-positive puncta (Fig. 4 B), providing another indication of altered intracellular organization. The molecular identity of these puncta and their relationship to centrosome disruption or changes in microtubule organization will require further study, however.

In summary, our findings identify NDE1 as a spatially defined subdistal appendage-associated factor occupying an intermediate domain between the inner mother-centriole architecture and the more peripheral microtubule- and motor-associated environment. NDE1 contributes to centrosome integrity and microtubule-organizing activity, while its depletion is accompanied by broader changes in centriolar and autophagy-associated compartments. Together, these results support a view of subdistal appendages as modular platforms through which the mature mother centriole coordinates centrosome architecture, microtubule organization, and intracellular trafficking.

## Acknowledgements

We are deeply grateful to Dr. Meng-Fu Bryan Tsou and Dr. Kanako Ozaki for generously teaching us ExSTED and for sharing important cell-line resources that made this work possible. We thank Dr. David Prober, Dr. Marianne Bronner, Dr. Rebecca Maria Voorhees, and Dr. Akiko Kumagai for their thoughtful discussions, advice, and encouragement. We also thank Biological Imaging Facility and Flow Cytometry and Cell Sorting Facility at Caltech, as well as Dr. Mary Grace Velasco and Kristofer Fertig from Abberior, for their support with imaging. We are sincerely grateful to all members of the Glover laboratory for creating a supportive environment. We also thank Dr. Inês Baião dos Santos for her generous help and kindness. This work was supported in large part by the National Institutes of Health under award number R01CA259382.

## Methods

### Cell culture and transfection

MEFs were immortalized as previously described^27^ and cultured in DMEM-Glutamax (GIBCO) with 10% fetal bovine serum (FBS, VWR) and penicillin-streptomycin. RPE-1 cells were provided by Dr. Meng-Fu Bryan Tsou and Dr. Kanako Ozaki at MSK. RPE-1 cells were cultured in DMEM/F12 (GIBCO) with 10% fetal bovine serum (VWR), 2 mM Glutamax (GIBCO) and penicillin-streptomycin. All cells were cultured at 37 °C in an atmosphere containing 5% CO2. Cells were routinely tested for mycoplasma contamination, with all experiments performed on mycoplasma-negative cultures. For knockdown experiments, all cells were transfected by Lipofectamine RNAi MAX (Invitrogen) according to the manufacturer’s instructions. In a 24-well format, each transfection mixture consisted of 6 pmol siRNA duplex and 1 µl Lipofectamine RNAi MAX in 100 µl Opti-MEM (GIBCO). The diluted RNAi molecules were thoroughly mixed and incubated for 20 minutes at room temperature. MEFs were then seeded at 2.5 × 10^4^ cells in 500 µL of DMEM complete growth medium and mixed with the transfection mixture. Cells were counted and fixed 96 h after transfection. The NDE1 siRNA sequence was confirmed to have no sequence complementarity to NDEL1. To generate MEF cell lines stably overexpressing hNDE1, lentivirus was produced in HEK293T cells by co-transfection of the hNDE1 lentiviral expression plasmid with the packaging plasmid psPAX2 and the envelope plasmid pMD2.G. MEFs were then transduced with the resulting lentivirus in the presence of LentiBOOST™ Transduction Enhancer (BioLegend/Revvity), according to the manufacturer’s instructions. Transduced cells were selected with blasticidin for 7 days. hNDE1 expression was then verified by qPCR and Western blotting.

### Immunoblotting

Unless otherwise stated, flash-frozen cell pellets were lysed directly in 4x Laemmli Sample buffer (BioRad) supplemented with 10% β-mercaptoethanol. Equal protein amounts, estimated from cell number, were subjected to electrophoresis on 10% or 15% Tris-Glycine Acrylamide (BioRad, 30% Acrylamide/Bis 29:1) gels, transferred to nitrocellulose (BioRad) and immunoblotted with the indicated primary and secondary antibodies. Data analysis was performed with Image Lab (BioRad).

### Quantification of autophagic flux using LC3B and p62 Western blotting

Cells were treated with bafilomycin A1 (BafA1; Cell Signaling Technology, #50-205-0494) at 0.1 µM for either 4 or 6 hours. Following treatment, cells were washed with ice-cold PBS and lysed in RIPA buffer containing protease and phosphatase inhibitors. Western blotting was conducted, and membranes were sequentially incubated with antibodies targeting LC3B, p62, and GAPDH. Band intensities were quantified from ChemiDoc MP images using Image Lab v6.1.0 (BioRad). LC3B-II and p62 levels were normalized to GAPDH detected on the same membranes. Autophagic flux was quantified based on the BafA1-dependent accumulation of LC3B-II relative to the corresponding untreated condition, with p62 serving as an additional indicator of autophagic cargo accumulation. Quantifications were derived from at least three independent biological replicates. Individual data points are provided in the Supplementary Data referenced for each figure.

### Immunofluorescence

Cells were grown on coverslips, washed with 1x PBS after removal of the media, and then fixed in cold methanol at -20°C for at least 12 minutes. Cells were rehydrated in 1x PBS, permeabilized in PBS containing 0.5% Triton X-100 for 15 minutes, and washed three times for 10 minutes in PBS containing 0.1% Triton X-100. Blocking was performed in PBS containing 0.1% Tween-20, 10% FBS for 1h, followed by incubation with primary and secondary antibodies in PBS containing 0.1% Tween-20, 10% FBS at 4°C overnight respectively. Washes were performed using PBS containing 0.1% Tween-20. Coverslips were mounted in Vectashield Mounting Medium (Vector Laboratories, H1200-10). For STED, coverslips were mounted in Abberior Mount Solid Antifade.

Confocal images were acquired using a Leica Stellaris/SP8 confocal microscope equipped with a 63×/1.4 NA oil-immersion objective and controlled with Leica Application Suite X software (LAS X). Images were deconvolved using Huygens Professional version 19.04 (Scientific Volume Imaging, The Netherlands). STED images, together with corresponding confocal images, were acquired on an Abberior STEDYCON system using a 100× oil-immersion objective and deconvolved using Abberior LIGHTBOX. Image processing and analysis were performed in ImageJ/Fiji and CellProfiler. Images shown represent projections of all acquired z-sections, collected with a z-step size of ≤0.5 µm.

### Expansion Microscopy

Expansion microscopy used in this work was adapted from protocol provided by Dr. Meng-Fu Bryan Tsou and Dr. Kanako Ozaki at MSK. Cells were fixed in ice-cold methanol at −20°C for 10 minutes, infused with 2% acrylamide and 1.4% formaldehyde in PBS for 5 h at 37°C, and embedded in expansion gel polymerized with APS and TEMED for 1 h at 37°C. Gels were denatured at 95°C for 90 minutes, washed, expanded in deionized water, and then immunostained with the indicated primary and secondary antibodies in PBS-based blocking buffer containing 10% FBS and 0.1% Tween-20. After final expansion in deionized water, gels were mounted cell-side down on glass-bottom dishes for imaging.

### Graphical Analysis and Statistics

All data represent at least three independent experiments unless otherwise indicated. Statistical significance for comparisons with control conditions was determined using two-tailed unpaired t tests unless otherwise indicated. Exact sample sizes, statistical tests, and significance values are indicated in the corresponding figure legends. All analyses were performed with GraphPad Prism 10.

### qRT-PCR analysis

RNA samples were prepared using the Qiagen RNeasy Micro Kit (#74004). qRT-PCR was performed using the Luna Universal One-Step RT-qPCR Kit (New England Biolabs) according to the manufacturer’s instructions. Amplification was monitored with SYBR Green chemistry on an Applied Biosystems StepOnePlus Real-Time PCR System. Each reaction was run in triplicate. Gene expression levels were normalized to GAPDH as a housekeeping gene, and relative expression was calculated using the ΔΔCt method.

## RESOURCE AVAILABILITY

### Lead contact

Requests for further information and resources should be directed to and will be fulfilled by the lead contact, David M. Glover.

### Materials availability

Plasmids and other materials used in this study are available from the lead contact upon reasonable request, subject to any restrictions associated with their original sources.

### Data and code availability

Quantitative source data are provided with this paper. Custom image-analysis scripts are available from the lead contact upon reasonable request.

## AUTHOR CONTRIBUTIONS

V.Y., P.A.C., and D.M.G. conceived and designed the project. V.Y. performed the majority of the experiments, analyzed the data, and wrote the initial draft of the manuscript. P.A.C. established the early stages of the project, contributed to the autophagy experiments, and provided critical feedback, suggestions, and discussion throughout the study. S.M.S. contributed to the microscopy-based analysis of centrosome number. V.Y., P.A.C., and D.M.G. critically revised and edited the manuscript.

---

**Figure S1. Additional characterization of NDE1 localization to the mother centriole subdistal appendage region in MEFs and RPE-1 cells. (A)** Additional deconvolved ExM-confocal and ExSTED images showing NDE1 localization at the appendage-bearing region of the mother centriole during interphase in MEFs. NDE1 signal is also detected near the proximal ends of the daughter centriole. The centriole cylinder is marked by acetylated tubulin (Ac-tub, cyan), and NDE1 is shown in magenta. Schematics at the upper right indicate the corresponding centriole orientation. Scale bars, 1 µm. **(B)** Deconvolved ExM-confocal and ExSTED images showing NDE1 localization at the appendage-bearing region of the mother centriole during interphase in RPE-1 cells. Representative G1 and S/G2 centrosomes are shown in panels i and ii, respectively. The centriole cylinder is marked by acetylated tubulin (Ac-tub, cyan), and NDE1 is shown in magenta. During mitosis, NDE1 staining at the appendage region is reduced (iii), and NDE1 is also detected at the cytokinetic ring/bridge (iv). Schematics below the images indicate the corresponding centriole orientation. Scale bars, 0.5 µm in i-iii and 1 µm in iv. **(C)** Additional ExM-confocal and ExSTED images showing that NDE1 localizes in close proximity to CEP170 at the mother centriole in RPE-1 cells. NDE1 forms a ring-like pattern around the Ac-tub-labeled centriole cylinder. Schematics below the images indicate the corresponding centriole orientation. Scale bar, 1 µm. In some examples, NDE1 and CEP170 signal are also detected near the daughter centriole, potentially corresponding to a proximal centriolar pool. **(D)** Quantification of ring diameter from ExSTED images in RPE-1 cells. Schematics indicate the corresponding centriole orientation. Each point represents one mother centriole. Black bars indicate the mean with 95% confidence interval. Exact sample sizes: ODF2, n = 10; CEP128, n = 11; NDE1, n = 10; NIN, n = 10; and CEP170, n = 13 from 3 independent experiments. **(E)** Quantification of the axial position of each signal relative to the Ac-tub-marked centriole base from ExSTED images in RPE-1 cells. Schematics indicate the corresponding centriole orientation. Each point represents one mother centriole measurement. Black bars indicate the median with interquartile range. Exact sample sizes: ODF2, n = 22; CEP128, n = 14; NDE1, n = 12; and NIN, n = 10 from 3 independent experiments.

**Figure S2. Additional analysis of ODF2- and CEP128-dependent NDE1 centrosomal localization in RPE-1 cells. (A)** Representative ExSTED images of NDE1 staining in control, ODF2-depleted and CEP128-depleted RPE-1 cells. The centriole cylinder is marked by acetylated tubulin (Ac-tub, cyan), and NDE1 is shown in magenta. Scale bar, 1 µm. **(B)** Quantification of centrosomal NDE1 fluorescence intensity in the indicated conditions from ExSTED images. Each point represents one mother centriole. Black bars indicate mean with 95% confidence interval. Statistical significance was determined by one-way ANOVA followed by Dunnett’s multiple-comparisons test comparing each depletion condition with the control; ****P < 0.0001; Exact sample sizes: Control, n = 10; ODF2 KD, n = 9; and CEP128 KD, n = 16 from 3 independent experiments. **(C)** Two-dimensional spatial mapping of ODF2, CEP128, NDE1, and NIN within the subdistal appendage in RPE-1 cells. Mean protein positions are plotted according to the measured diameter (x axis) and distance from the bottom of the mother centriole (y axis). Horizontal and vertical error bars indicate SD. The overlaid schematic depicts the approximate lateral geometry of the subdistal appendage and its relationship to the centriole.

**Figure S3. Additional characterization of centriole defects following NDE1 depletion in MEFs and RPE-1 cells. (A)** Additional ExSTED images showing reduced CPAP staining in NDE1-depleted MEFs. NDE1 depletion is associated with increased distance between paired centrioles and abnormal centriole morphology. The centriole cylinder is marked by acetylated tubulin (Ac-tub, cyan), and CPAP is shown in magenta. Scale bars, 0.5 µm. **(B)** Confocal images of control and NDE1-depleted MEFs stained for CEP152 and PLK4. CEP152 is shown in red, PLK4 in green, and nuclei are stained with DAPI in blue. NDE1 depletion is associated with increased distance between paired centrioles and reduced CEP152 staining. Scale bars, 3 µm. **(C)** Confocal images of control and NDE1-depleted RPE-1 cells stained for CEP152 and PLK4. CEP152 is shown in red, PLK4 in green, and nuclei are stained with DAPI in blue. NDE1 depletion is associated with increased distance between paired centrioles and reduced CEP152 staining. Scale bars, 3 µm. **(D)** Additional ExM-confocal images showing abnormal centriole morphology in NDE1-depleted MEFs. NDE1 depletion is associated with altered centriole organization and increased distance between paired centrioles. The centriole cylinder is marked by Ac-tub in red; distal centriolar markers CP110 or CEP164 are shown in yellow; and PLK4 is shown in green. Scale bars, 0.4 µm.

**Figure S4. Additional analysis of PCM organization and centrosome-focused microtubule growth following NDE1 depletion in MEFs and RPE-1 cells. (A)** Representative confocal images showing reduced centrosome-focused microtubule organization in interphase NDE1-depleted RPE-1 cells. Microtubules are shown in gray, CEP152 is shown in red, and nuclei are stained with DAPI in blue. Centrioles are defined by CEP152-positive foci. Scale bars, 3 µm. **(B)** Quantification of centriolar γ-tubulin fluorescence intensity in control, mNDE1-depleted, and mNDE1-depleted MEFs overexpressing hNDE1. Each dot represents the γ-tubulin fluorescence intensity associated with an individual centrosome. Black bars indicate the mean with 95% confidence interval. Statistical significance was determined by two-tailed unpaired t test for each comparison with the control condition; ****P < 0.0001, ns, not significant. Exact sample sizes: Control, n = 14; mNDE1 KD, n = 19; and mNDE1 KD+hNDE1 OE, n = 22 from 3 independent experiments. **(C)** Quantification of centriolar γ-tubulin fluorescence intensity in control and NDE1-depleted RPE-1 cells. Each dot represents the γ-tubulin fluorescence intensity associated with an individual centrosome. Black bars indicate the mean with 95% confidence interval. Statistical significance was determined by two-tailed unpaired t test; ns, not significant. Exact sample sizes: Control, n = 9; and NDE1 KD, n = 13 from 3 independent experiments. **(D)** Representative confocal images showing reduced centrosome-focused microtubule asters in interphase NDE1-depleted RPE-1 cells. EB1-positive microtubule plus ends are shown in yellow, and DNA is stained with DAPI in blue. Scale bars, 2 µm. **(E)** Quantification of centrosome-associated EB1 fluorescence intensity during interphase in control and NDE1-depleted RPE-1 cells. Each dot represents the EB1 fluorescence intensity associated with an individual centrosome. Black bars indicate the mean with 95% confidence interval. Statistical significance was determined by two-tailed unpaired t test for comparison with the control condition; ***P < 0.001. Exact sample sizes: Control, n = 18; and NDE1 KD, n = 19 from 3 independent experiments. **(F)** Quantification of centrosome-associated pericentrin fluorescence intensity during interphase in control and NDE1-depleted RPE-1 cells. Each dot represents the pericentrin fluorescence intensity associated with an individual centrosome. Black bars indicate the mean with 95% confidence interval. Statistical significance was determined by two-tailed unpaired t test for comparison with the control condition; ****P < 0.0001. Exact sample sizes: Control, n = 19; and NDE1 KD, n = 19 from 3 independent experiments. **(G)** Representative confocal images illustrating microtubule regrowth around centrosomes in MEFs at 0, 2, 10, and 60 minutes after nocodazole washout. Pericentrin, which marks the centrosome, is shown in green; EB1-positive microtubule plus ends are shown in yellow; α-tubulin is shown in red; and DNA is stained with DAPI in blue. Scale bars, 5 µm.

**Figure S5. NDE1 depletion perturbs autophagic compartment homeostasis. (A)** Autophagic flux was assessed by immunoblot analysis of the phagophore- and autophagosome-associated lipidated form of LC3B (LC3B-II) in control and NDE1-depleted MEFs treated with or without bafilomycin A1 (BafA1) for 4 or 6 h. GAPDH was used as a loading control. **(B)** Densitometric quantification of LC3B-II band intensity normalized to GAPDH in control and NDE1-depleted MEFs without BafA1. Black bars indicate the mean with 95% confidence interval. Statistical significance was determined by two-tailed unpaired t test for comparison with the control condition; ns, not significant. Exact sample sizes: Control, n = 5; and NDE1 KD, n = 5 from 5 independent experiments. **(C)** Autophagic flux was calculated as the difference between normalized LC3B-II levels in BafA1-treated cells and the corresponding untreated cells at 4 and 6 h. Data are presented as mean with 95% confidence interval. Statistical significance was determined using two-tailed unpaired t test. Exact sample sizes: Control, n = 5; and NDE1 KD, n = 5 from 5 independent experiments. **(D)** Representative immunoblot analysis of SQSTM1/p62 in control and NDE1-depleted MEFs treated with or without BafA1 for 4 or 6 h. GAPDH was used as a loading control. **(E)** Densitometric quantification of SQSTM1/p62 band intensity normalized to GAPDH on the same membranes. Values are presented as fold change relative to the corresponding control and as mean ± SEM from three independent biological replicates. Statistical significance was determined using two-tailed unpaired t test for comparison with the corresponding control condition. With or without BafA1 for 4 or 6 h, exact sample sizes: Control, n = 3; and NDE1 KD, n = 3 from 3 independent experiments. **(F)** Representative confocal images showing increased LC3B signal in NDE1-depleted MEFs. LC3B is shown in red, and nuclei are stained with DAPI in blue. Boxed regions indicate areas shown at higher magnification. Scale bars, 10 µm (4 µm in magnified views). **(G)** Quantification of LC3B fluorescence intensity in control, mNDE1-depleted, and mNDE1-depleted MEFs overexpressing hNDE1. Black bars indicate the mean with 95% confidence interval. Statistical significance was determined by two-tailed unpaired t test for each comparison with the control condition; ***P < 0.001, ns, not significant. Exact sample sizes: Control, n = 34; mNDE1 KD, n = 38; and mNDE1 KD+hNDE1 OE, n = 45 from 3 independent experiments. **(H-I)** Quantification of autophagic compartments using the tandem fluorescent LC3B reporter. (H) Autophagosome number per unit cell area was increased following NDE1 depletion, whereas (I) autolysosome number per unit cell area was not significantly altered. Values are normalized as indicated. Black bars indicate the mean with 95% confidence interval. Statistical significance was determined by two-tailed unpaired t test for each comparison with the control condition; **P < 0.01, ns, not significant. For both analyses, exact sample sizes: Control, n = 52; and NDE1 KD, n = 46 from 3 independent experiments.

**Figure S6. Additional analysis of autophagic compartments following NDE1 depletion. (A)** Representative immunoblot analysis of LC3B in control and NDE1-depleted RPE-1 cells treated with or without BafA1 for 1.5 or 3 h. LC3B-I and LC3B-II are indicated; GAPDH was used as a loading control. **(B)** Quantification of basal LC3B-II levels normalized to GAPDH in control and NDE1-depleted RPE-1 cells. Black bars indicate the mean with 95% confidence interval. Statistical significance was determined by two-tailed unpaired t test for comparison with the control condition; ns, not significant. Exact sample sizes: Control, n = 7; and NDE1 KD, n = 7 from 7 independent experiments. **(C)** Quantification of autophagic flux following 1.5- or 3-h BafA1 treatment. Black bars indicate the mean with 95% confidence interval. Statistical significance was determined by two-tailed unpaired t test for comparison with the control condition; ns, not significant. Exact sample sizes: Control, n = 7; and NDE1 KD, n = 7 from 7 independent experiments. **(D)** Representative confocal images showing increased p62 puncta in NDE1-depleted MEFs. p62 is shown in red, and nuclei are stained with DAPI in blue. Scale bars, 10 µm. **(E)** Representative immunoblot analysis of SQSTM1/p62 in control and NDE1-depleted RPE-1 cells. GAPDH was used as a loading control. **(F)** Densitometric quantification of SQSTM1/p62 band intensity normalized to GAPDH. Values are presented as fold change relative to the corresponding control and as mean ± 95% confidence interval. Statistical significance was determined using two-tailed unpaired t test for comparison with the corresponding control condition. Exact sample sizes: Control, n = 3; and NDE1 KD, n = 3 from 3 independent experiments. **(G)** Representative confocal images showing increased p62 puncta in NDE1-depleted RPE-1 cells. p62 is shown in green, and nuclei are stained with DAPI in blue. Scale bars, 10 µm. **(H)** Representative confocal images showing LC3B signal in control and NDE1-depleted RPE-1 cells treated with or without bafilomycin A1 for 24 h. LC3B is shown in red, and nuclei are stained with DAPI in blue. Scale bars, 10 µm. **(I)** Quantification of mCherry-LAMP1 puncta number per unit cell area in control and NDE1-depleted MEFs. Each point represents one individual cell. Black bars indicate the mean with 95% confidence interval. Statistical significance was determined by two-tailed unpaired t test; ns, not significant; Exact sample sizes: Control, n = 74; and NDE1 KD, n = 57 from 3 independent experiments.

**Figure S7. Proposed model for altered autophagic compartment dynamics following NDE1 depletion.** Schematic illustrating the major stages of autophagy and summarizing the autophagic phenotype associated with NDE1 depletion. During autophagosome formation, cytoplasmic LC3B-I is lipidated to generate membrane-associated LC3B-II, while p62/SQSTM1 associates with cargo destined for degradation. Mature autophagosomes subsequently fuse with LAMP1-positive lysosomes to form autolysosomes, where cargo and p62 are degraded and LC3 components are recycled. NDE1 depletion is associated with an increased LC3B-II pool and accumulation of LC3B-positive autophagosomes, whereas the abundance of autolysosomes is comparatively unchanged. Together with the preservation of ongoing autophagic flux, these findings are consistent with an expansion of the autophagosome pool and altered balance between autophagic compartments rather than a complete block in autophagosome-lysosome fusion or autolysosome formation.

