## Supplementary material for "NDE1 Localizes to the Subdistal Appendages to Maintain Centrosome Integrity and Microtubule Organization": Supp Fig 1

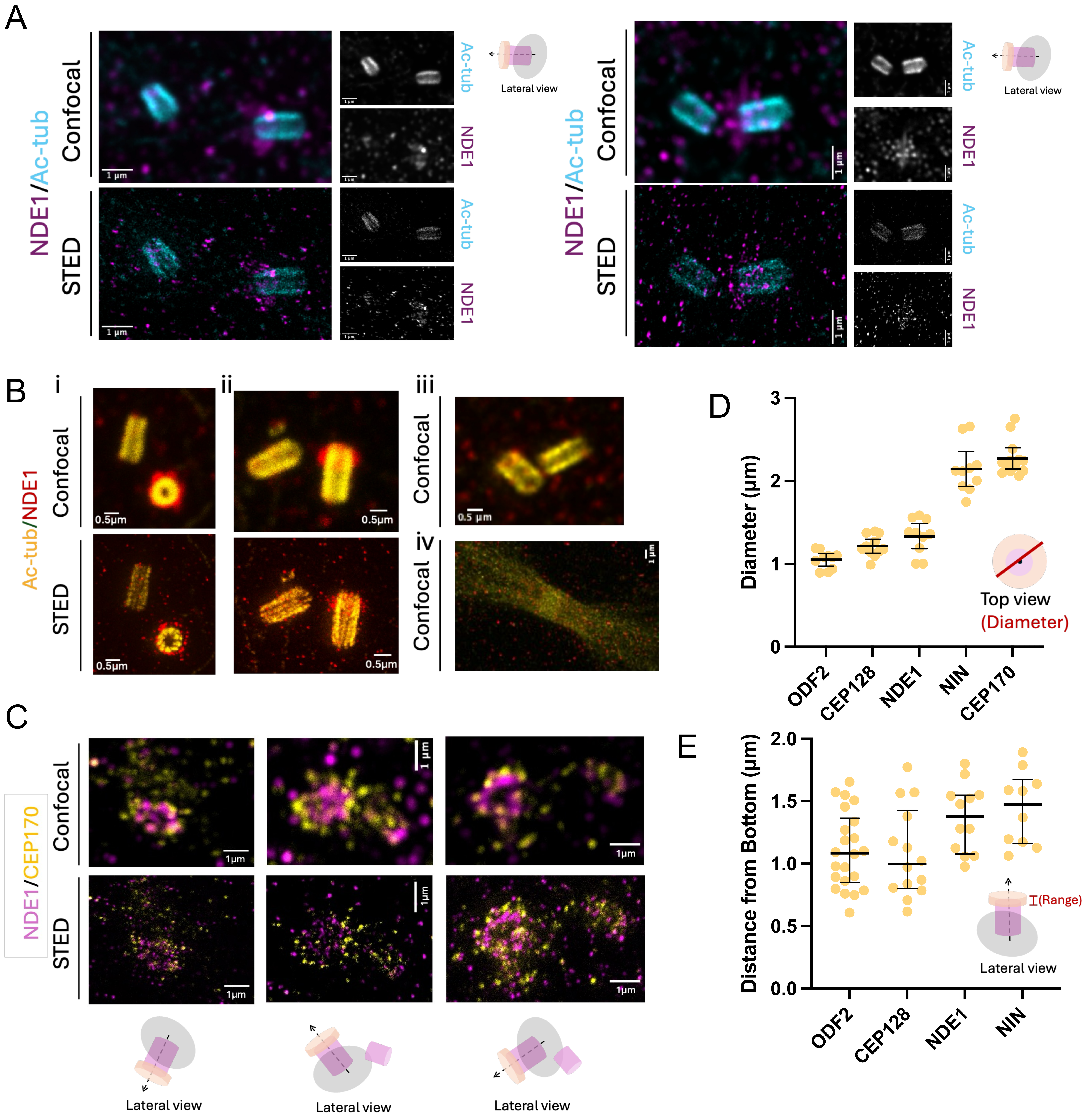

A
NDE1/Ac-tub
Confocal
STED
1 μm
Ac-tub
NDE1
Lateral view
B
i
ii
iii
iv
Ac-tub/NDE1
Confocal
STED
0.5μm
0.5 μm
1 μm
C
NDE1/CEP170
Confocal
STED
1μm
1 μm
Lateral view
D
Diameter (μm)
0
1
2
3
ODF2
CEP128
NDE1
NIN
CEP170
Top view
(Diameter)
E
Distance from Bottom (μm)
0.0
0.5
1.0
1.5
2.0
ODF2
CEP128
NDE1
NIN
(Range)
Lateral view
