## Supplementary material for "NDE1 Localizes to the Subdistal Appendages to Maintain Centrosome Integrity and Microtubule Organization": Supp Fig 2

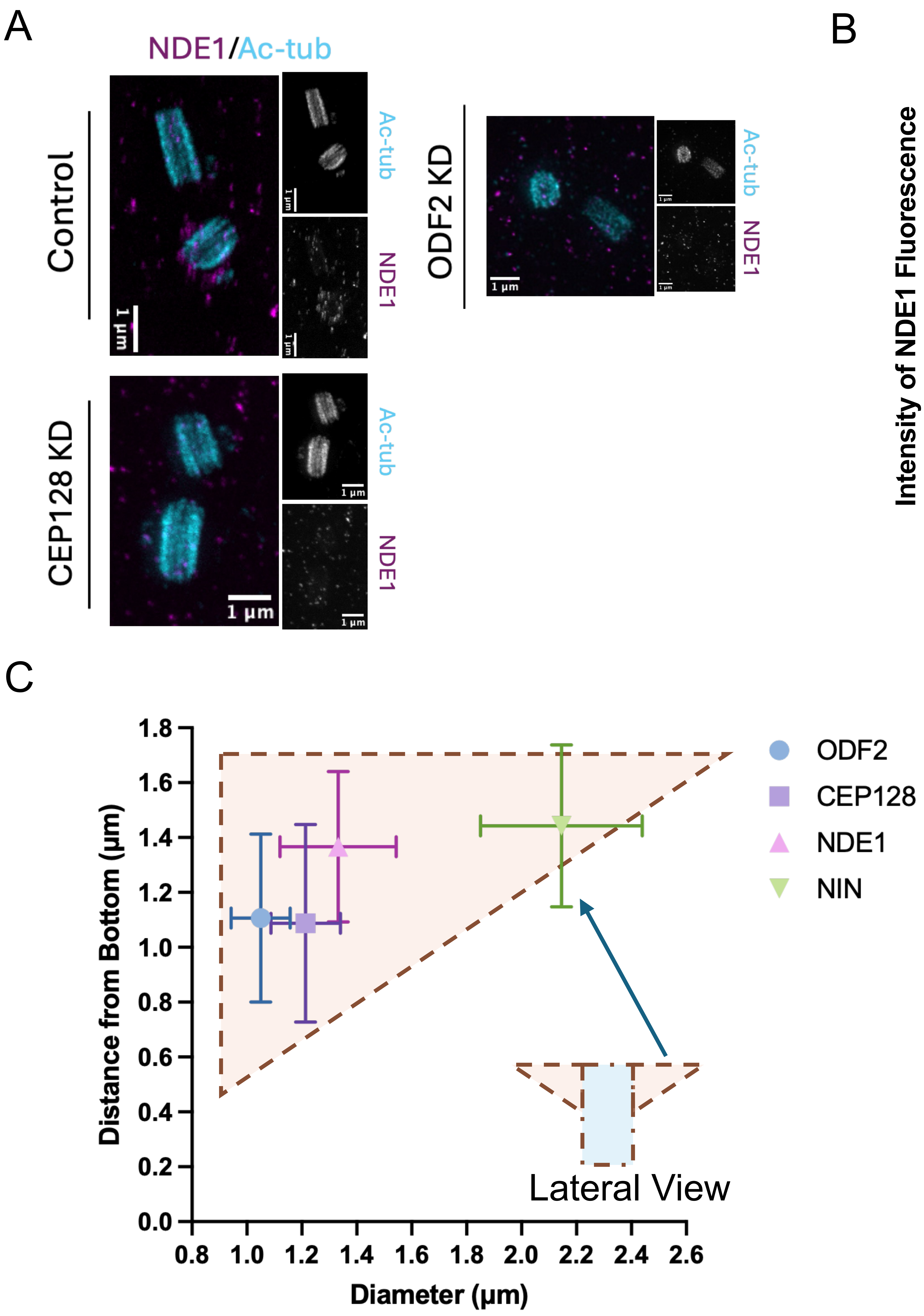
A
NDE1/Ac-tub
Control
CEP128 KD
ODF2 KD
Ac-tub
NDE1
1 µm
B
Intensity of NDE1 Fluorescence Signal (Normalized)
1.2
0.9
0.6
0.3
0.0
****
****
Control
CEP128 KD
ODF2 KD
C
Distance from Bottom (µm)
1.8
1.6
1.4
1.2
1.0
0.8
0.6
0.4
0.2
0.0
0.8
1.0
1.2
1.4
1.6
1.8
2.0
2.2
2.4
2.6
Diameter (µm)
Lateral View
ODF2
CEP128
NDE1
NIN
