## Supplementary material for "NDE1 Localizes to the Subdistal Appendages to Maintain Centrosome Integrity and Microtubule Organization": Supp Fig 3

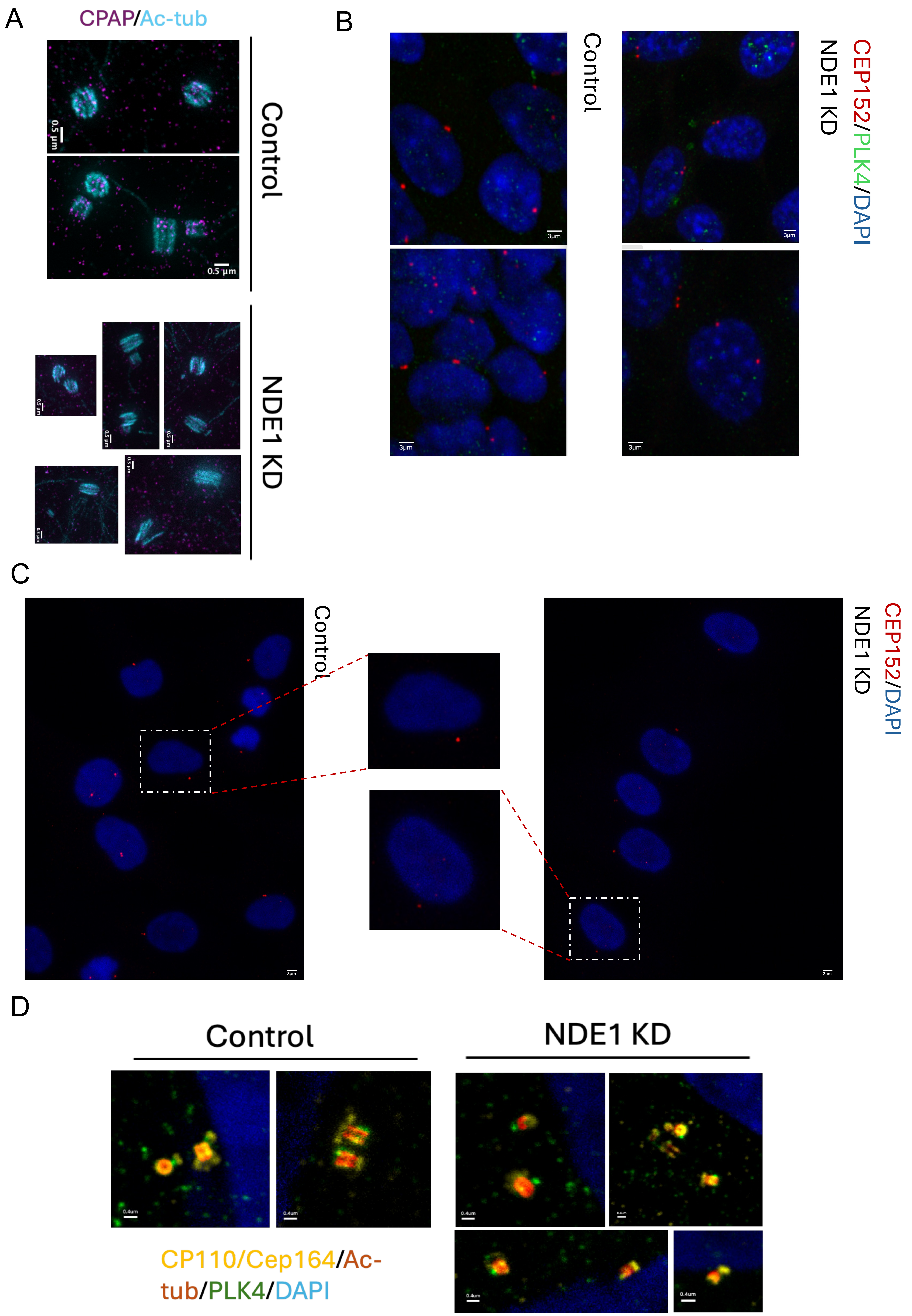
A
CPAP/Ac-tub
0.5 µm
Control
NDE1 KD
B
Control
NDE1 KD
CEP152/PLK4/DAPI
3µm
C
Control
NDE1 KD
CEP152/DAPI
D
Control
NDE1 KD
0.4um
CP110/Cep164/Ac-tub/PLK4/DAPI
