## Supplementary material for "NDE1 Localizes to the Subdistal Appendages to Maintain Centrosome Integrity and Microtubule Organization": Supp Fig 4

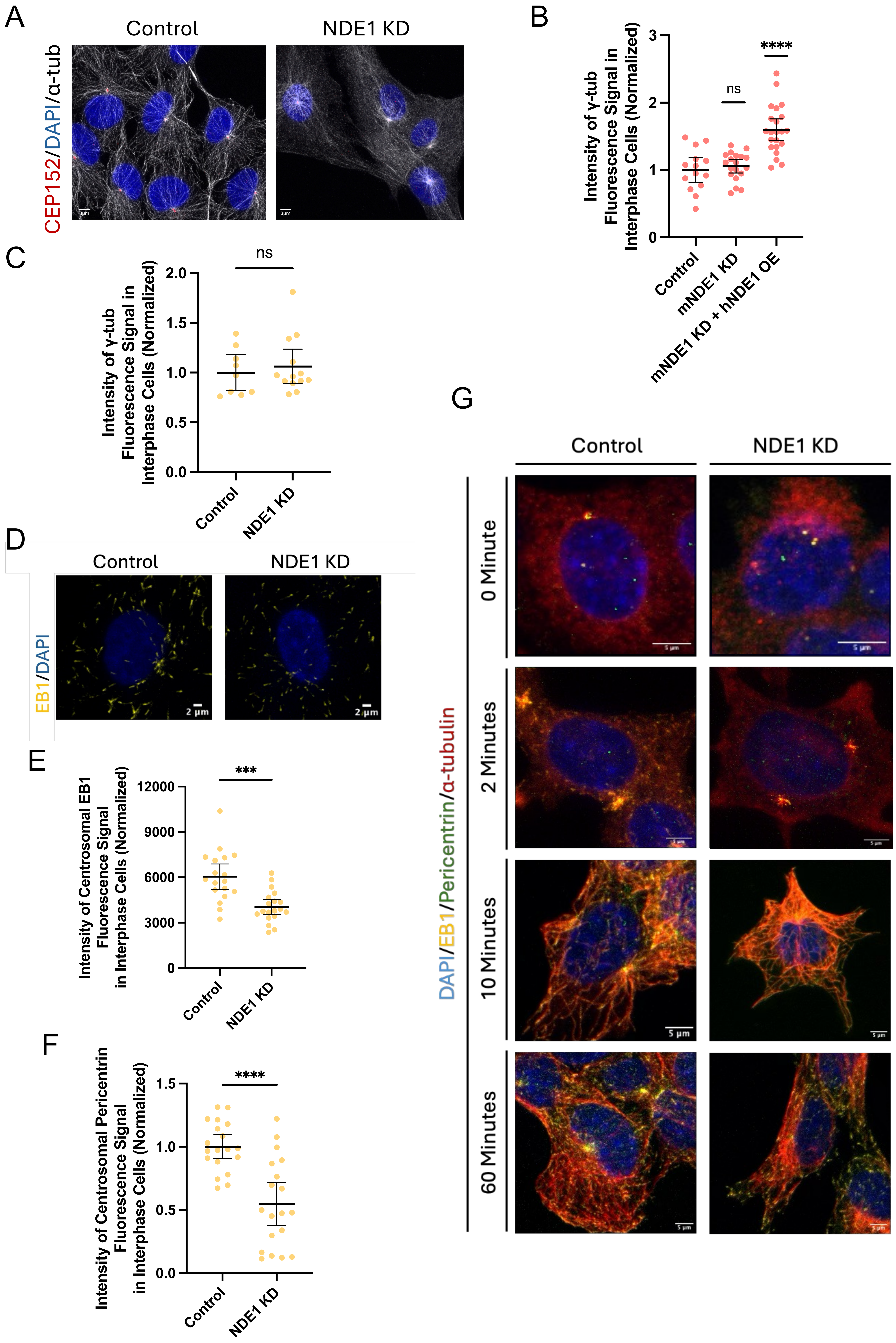

A
Control
NDE1 KD
CEP152/DAPI/α-tub
B
Intensity of γ-tub Fluorescence Signal in Interphase Cells (Normalized)
0
1
2
3
ns
****
Control
mNDE1 KD
mNDE1 KD + hNDE1 OE
C
Intensity of γ-tub Fluorescence Signal in Interphase Cells (Normalized)
0.0
0.5
1.0
1.5
2.0
ns
Control
NDE1 KD
D
Control
NDE1 KD
EB1/DAPI
2 μm
2 μm
E
Intensity of Centrosomal EB1 Fluorescence Signal in Interphase Cells (Normalized)
0
3000
6000
9000
12000
***
Control
NDE1 KD
F
Intensity of Centrosomal Pericentrin Fluorescence Signal in Interphase Cells (Normalized)
0.0
0.5
1.0
1.5
****
Control
NDE1 KD
G
Control
NDE1 KD
DAPI/EB1/Pericentrin/α-tubulin
0 Minute
2 Minutes
10 Minutes
60 Minutes
5 μm
