## Supplementary material for "NDE1 Localizes to the Subdistal Appendages to Maintain Centrosome Integrity and Microtubule Organization": Supp Fig 5

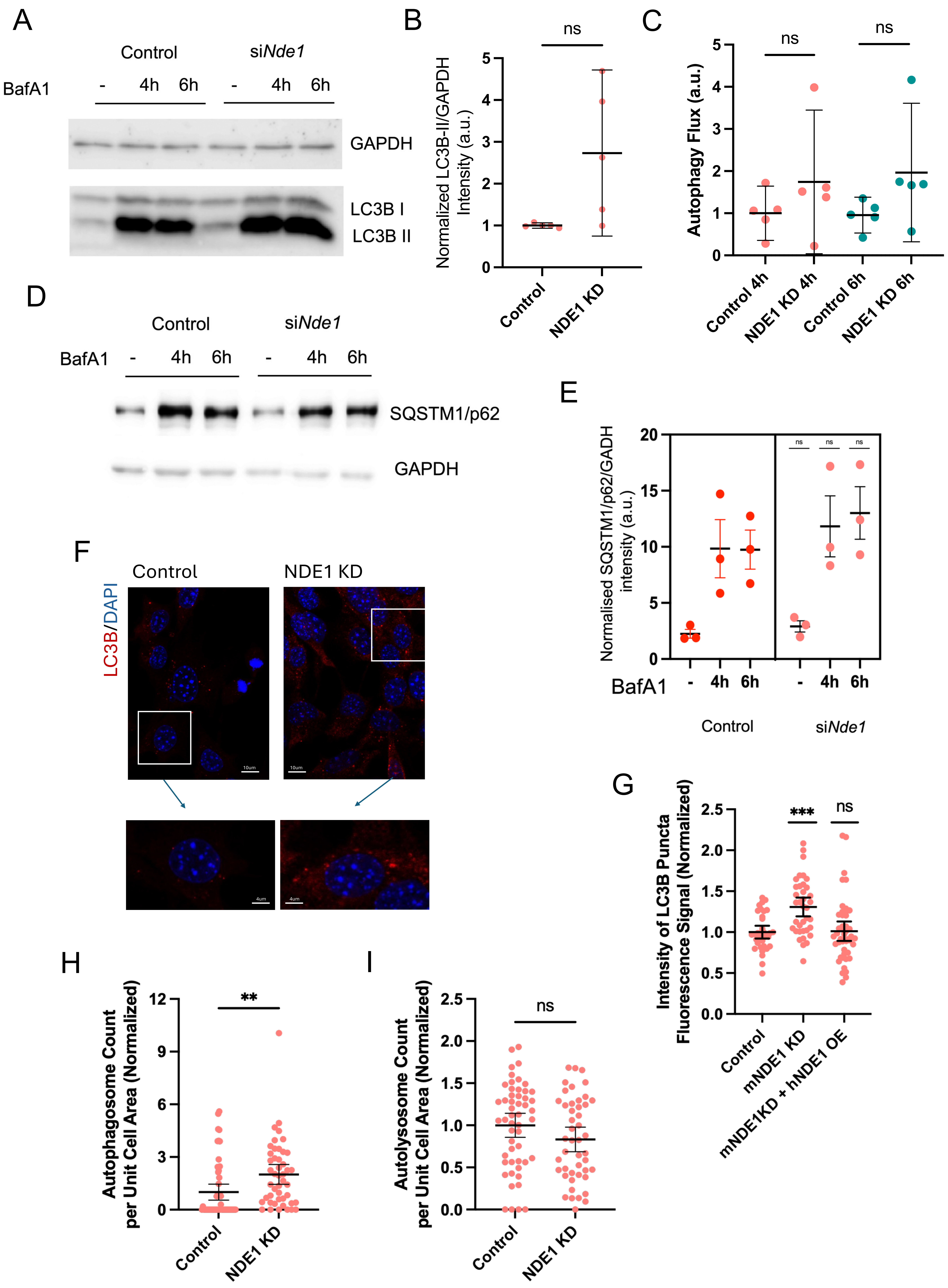

A
Control
siNde1
BafA1
- 4h 6h - 4h 6h
GAPDH
LC3B I
LC3B II
B
ns
Normalized LC3B-II/GAPDH Intensity (a.u.)
Control
NDE1 KD
C
ns
ns
Autophagy Flux (a.u.)
Control 4h
NDE1 KD 4h
Control 6h
NDE1 KD 6h
D
Control
siNde1
BafA1
- 4h 6h - 4h 6h
SQSTM1/p62
GAPDH
E
Normalised SQSTM1/p62/GADH intensity (a.u.)
ns ns ns
BafA1
- 4h 6h - 4h 6h
Control
siNde1
F
LC3B/DAPI
Control
NDE1 KD
G
Intensity of LC3B Puncta Fluorescence Signal (Normalized)
***
ns
Control
mNDE1 KD
mNDE1KD + hNDE1 OE
H
Autophagosome Count per Unit Cell Area (Normalized)
**
Control
NDE1 KD
I
Autolysosome Count per Unit Cell Area (Normalized)
ns
Control
NDE1 KD
