## Supplementary material for "NDE1 Localizes to the Subdistal Appendages to Maintain Centrosome Integrity and Microtubule Organization": Supp Fig 6

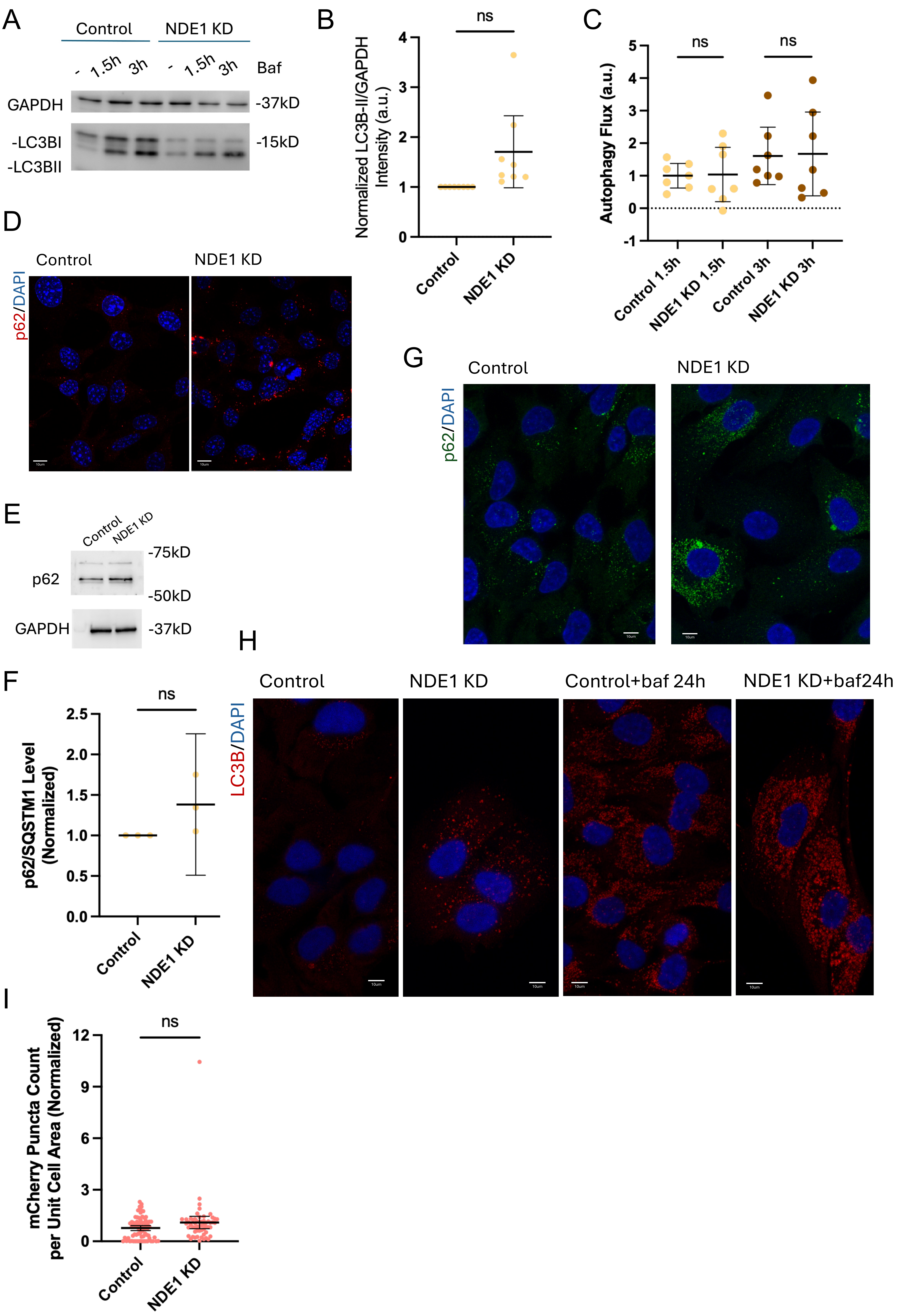
A
Control
NDE1 KD
- 1.5h 3h - 1.5h 3h Baf
GAPDH
-37kD
-LC3BI
-LC3BII
-15kD
B
ns
Normalized LC3B-II/GAPDH Intensity (a.u.)
0
1
2
3
4
Control
NDE1 KD
C
ns
ns
Autophagy Flux (a.u.)
-1
0
1
2
3
4
5
Control 1.5h
NDE1 KD 1.5h
Control 3h
NDE1 KD 3h
D
Control
NDE1 KD
p62/DAPI
E
Control
NDE1 KD
-75kD
p62
-50kD
GAPDH
-37kD
G
Control
NDE1 KD
p62/DAPI
F
ns
p62/SQSTM1 Level (Normalized)
0.0
0.5
1.0
1.5
2.0
2.5
Control
NDE1 KD
H
Control
NDE1 KD
Control+baf 24h
NDE1 KD+baf24h
LC3B/DAPI
I
ns
mCherry Puncta Count per Unit Cell Area (Normalized)
0
3
6
9
12
Control
NDE1 KD
