## Supplementary material for "NDE1 Localizes to the Subdistal Appendages to Maintain Centrosome Integrity and Microtubule Organization": Supp Fig 7

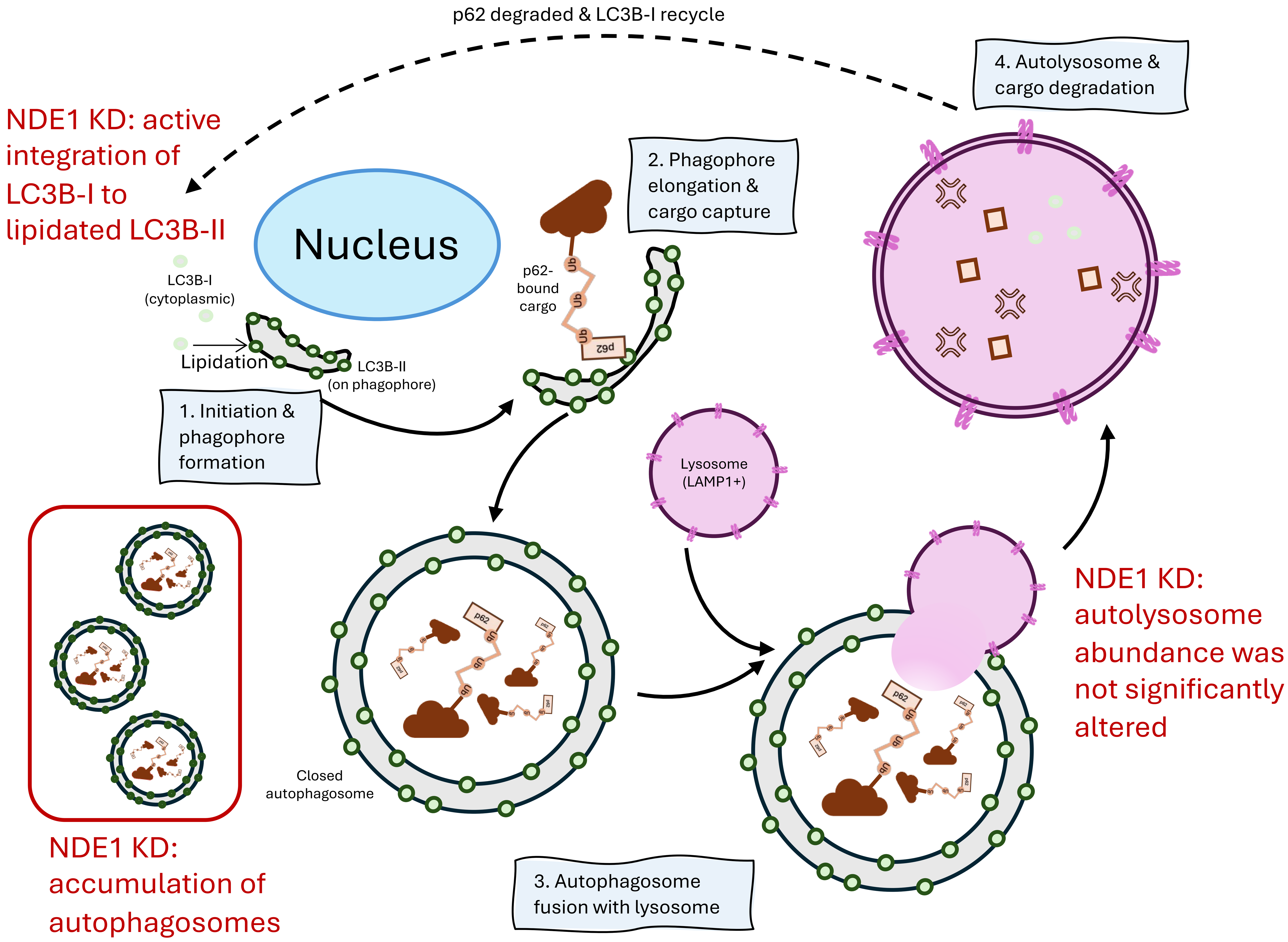
p62 degraded & LC3B-I recycle
4. Autolysosome & cargo degradation
NDE1 KD: active integration of LC3B-I to lipidated LC3B-II
Nucleus
2. Phagophore elongation & cargo capture
p62-bound cargo
LC3B-I (cytoplasmic)
Lipidation
LC3B-II (on phagophore)
1. Initiation & phagophore formation
Lysosome (LAMP1+)
NDE1 KD: autolysosome abundance was not significantly altered
Closed autophagosome
NDE1 KD: accumulation of autophagosomes
3. Autophagosome fusion with lysosome
